# Learning on a Limb: An outreach module to engage high school students in orthopaedics

**DOI:** 10.1101/2024.09.16.612729

**Authors:** Christopher J. Panebianco, Tala F. Azar, Michael P. Duffy, Madhura P. Nijsure, Emily Sharp, Margaret K. Tamburro, Michael Hast, Eileen M. Shore, Robert L. Mauck, Louis J. Soslowsky, Jamie R. Shuda, Sarah E. Gullbrand

## Abstract

Orthopaedic researchers need new strategies for engaging diverse students. Our field has demonstrated noticeable gaps in racial, ethnic, and gender diversity, which inhibit our ability to innovate and combat the severe socioeconomic burden of musculoskeletal disorders. Towards this goal, we designed, implemented, and evaluated Learning on a Limb, an orthopaedic research outreach module to teach diverse high school students about orthopaedic research. During the 4-hr module, students completed hands-on activities to learn how biomechanical testing, microcomputed tomography, cell culture, and histology are used in orthopaedic research. Over three years, we recruited 32 high school students from the Greater Philadelphia Area to participate in Learning on a Limb. Most participants identified as racial/ethnic or gender minorities in orthopaedic research. Using pre/post-tests, we found that students experienced significant learning gains of 51 percentage points from completing Learning on a Limb. In addition to teaching students about orthopaedic research, post-survey data demonstrated that participating in Learning on a Limb strongly influenced students’ interest in orthopaedic research. Several students acted on this interest by completing summer research experiences in the McKay Orthopaedic Research Laboratory at the University of Pennsylvania. Learning on a Limb instructors also benefited by having the opportunity to “pay it forward” to the next generation of students and build community within their department. Empowering institutions to host modules like Learning on a Limb would synergistically inspire diverse high school students and strengthen community within orthopaedic departments to ultimately enhance orthopaedic research innovations.

## 1.0 Introduction

Musculoskeletal disorders are a pressing public health concern. They are the leading cause of global disability and cost approximately $381 billion annually, making them the most expensive aggregated health category in the United States.^1,2^ To combat the high socioeconomic burdens of musculoskeletal disorders, orthopaedic researchers must continue studying the function, development, degeneration, and regeneration of orthopaedic tissues. Furthermore, we must engage students with orthopaedic research early in their education to galvanize them to advance the diagnosis, treatment, and prevention of musculoskeletal disorders.

K-12 outreach is effective for inspiring diverse students to pursue careers in science, technology, engineering, and mathematics (STEM).^3,4^ Such outreach modules are especially important for enhancing the representation of underrepresented minority (URM) students in STEM based on race, gender, and socioeconomic status.^5–7^ Increasing representation is important for combatting systemic barriers that URM populations have had to pursuing STEM careers and producing higher impact work.^8–10^ Thus, developing engaging K-12 outreach modules focused on orthopaedics will be crucial for diversifying the orthopaedics field and accelerating innovations in orthopaedic research.

The pedagogical approaches implemented in outreach modules are important for determining their efficacy. Traditional learning approaches, where instructors lecture students to teach information,^11^ produce non-active learners and promote superficial learning.^12,13^ Active learning is a favorable alternative, where instructors ask open-ended questions and use inquiry-led tasks to engage students in complex learning processes.^14,15^ Numerous studies demonstrated that implementing active learning techniques can broadly provide students with a deeper comprehension of the material being taught^16–18^ and reduce achievement gaps for URM students.^19–21^ Hands-on active learning is particularly effective for STEM outreach initiatives because it engages students and the physicality connects theoretical content to practical applications.^22^ Thus, we designed our orthopaedics outreach module using these pedagogies.

There are numerous published undergraduate bioengineering modules that are tangentially related to aspects of orthopaedic research explicitly; however, there are no modules that teach high school students about orthopaedic research. For example, modules have been designed to teach undergraduates about biomaterials,^23–26^ mechanobiology,^27–30^ microcomputed tomography (microCT)^31,32^, cell culture,^33–35^ and histology.^36,37^ Educators have successfully employed Next Generation Science Standards^38^ to adapt undergraduate modules for K-12 outreach;^39,40^ thus, we implemented similar standards to develop an orthopaedic outreach module for high school students.

Accordingly, we designed, implemented, and evaluated Learning on a Limb, an orthopaedic outreach module to teach diverse high school students about orthopaedic research. This module was designed as a half-day program for high school students in the Greater Philadelphia Area to learn about orthopaedic research using hands-on active learning. We implemented the module with three cohorts of high school students over three years and rigorously evaluated its efficacy using pre/post-tests and post-surveys.

## 2.0 Materials and methods

### 2.1 Module design and overview

Learning on a Limb was led by a diverse group of principal investigators, postdoctoral research fellows, graduate students, and research associates based in the McKay Orthopaedic Research Laboratory, which is affiliated with the Penn Center for Musculoskeletal Disorders (PCMD) and the Penn Achilles Tendinopathy Center of Research Translation. Approximately 6 months before the first session, volunteers began working with the Perelman School of Medicine’s Office of Outreach, Education, and Research (OER) to create a sustainable and impactful module, with learning goals that could be achieved using existing resources. Learning on a Limb was designed as a 4-hr module divided into three stages: (1) pre-activity exercises (∼35 min), (2) hands-on activities (∼2 hr and 35 min), and (3) post-activity exercises (∼50 min) (Figure 1).

**Figure 1.**
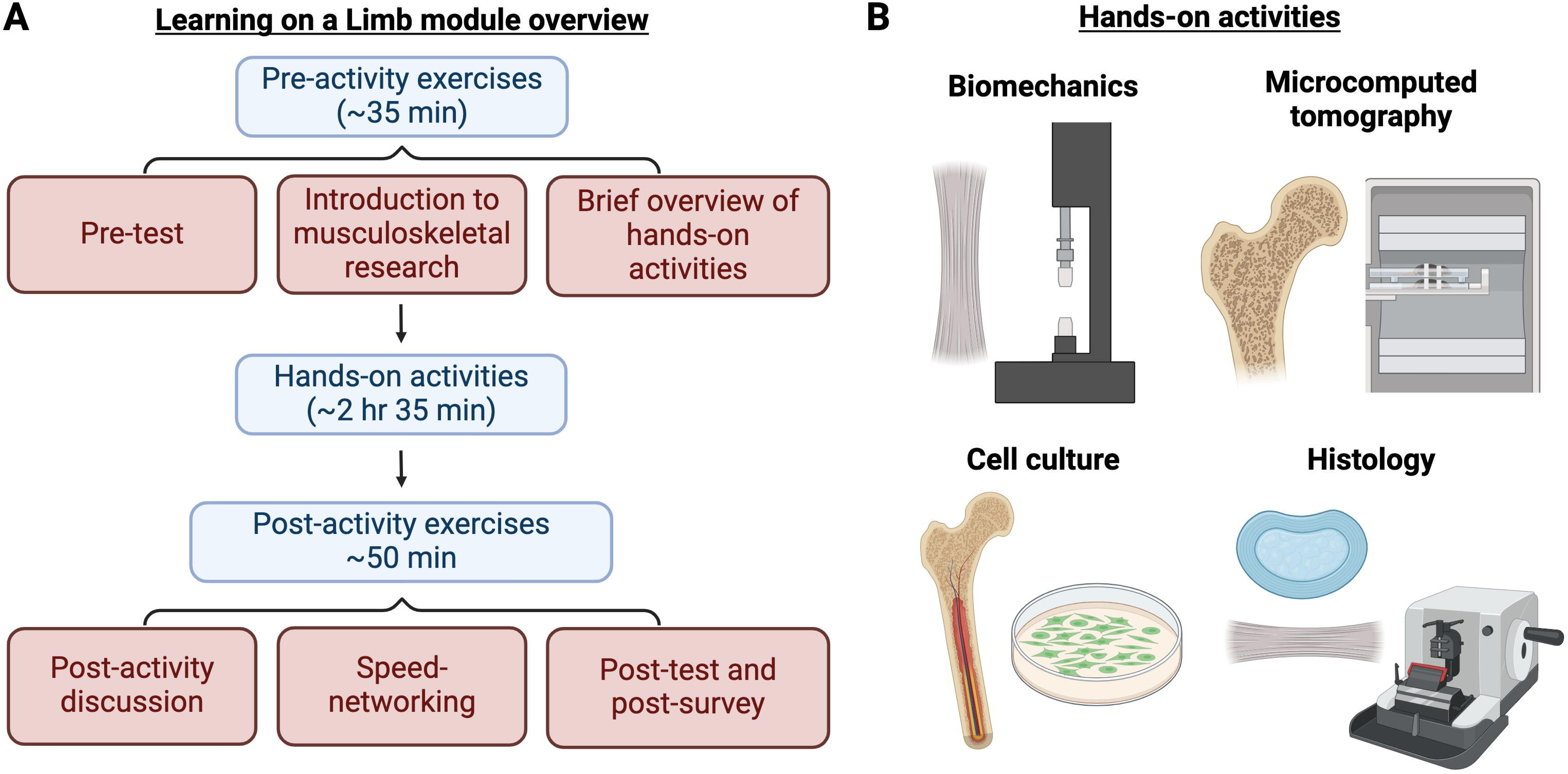
Learning on a Limb module overview. **(A)** Learning on a Limb was a 4-hr module consisting of pre-activity exercises, hands-on activities, and post-activity exercises. **(B)** High school students worked in groups of 2-4 to complete hands-on, circuit-style activities in biomechanical testing of rat tendons, microcomputed tomography (MicroCT) of rat bones, cell culture of mouse bone mesenchymal stromal cells (BMSCs), and histology of rat tendons and rabbit intervertebral discs (IVDs).

### 2.2 Student recruitment

Local high school students in the Greater Philadelphia Area were recruited using flyers sent through the Perelman School of Medicine Office of OER. We had 15 students in 2022, 12 students in 2023, and 5 students in 2024 participate in our study. Students worked in groups of 2-4 to complete all activities. All surveys were approved by the University of Pennsylvania Institutional Review Board (IRB Protocol # 823110 and 826159). Data was only analyzed for students who had a guardian complete an informed consent form.

### 2.3 Pre-activity exercises

During the pre-activity exercise, instructors provided background on orthopaedic research, techniques students would learn during the activities, and strategies for leveraging this experience to conduct future orthopaedic research (e.g., summer research experiences). We implemented active learning strategies throughout the pre-activity exercises (*e.g.,* multiple hands/voices)^41^ because active learning improves student performance in STEM courses and helps narrow achievement gaps experienced by underrepresented minority (URM) students.^19–21^

### 2.4 Biomechanical testing activity

The learning goals for the biomechanical testing activity were to understand basic principles of viscoelasticity and mechanical testing of orthopaedic tissues.

Prior to the module, Achilles tendons (one tendon per pair of students) from mature Sprague-Dawley rats were prepared, as described.^42^ Importantly, sex and weight of the rat were recorded at time of dissection to facilitate hypothesis generation during the module. Briefly, hindlimbs were separated, the ankle joint was disarticulated to free the muscle-tendon-foot complex, and all non-tendinous tissue removed (Figure 2A). Verhoeff’s stain was used to mark the tendon at the calcaneal insertion, 6 mm proximal to the insertion, and 8 mm proximal to the insertion. Sandpaper was adhered to the anterior and posterior surface of the tendon 8 mm proximal to the calcaneal insertion with cyanoacrylate, and the foot was potted in polymethyl methacrylate. Potted foot-tendon complexes were stored in 1X phosphate buffered saline (PBS) at 4°C until use (Figure 2B-C).

**Figure 2.**
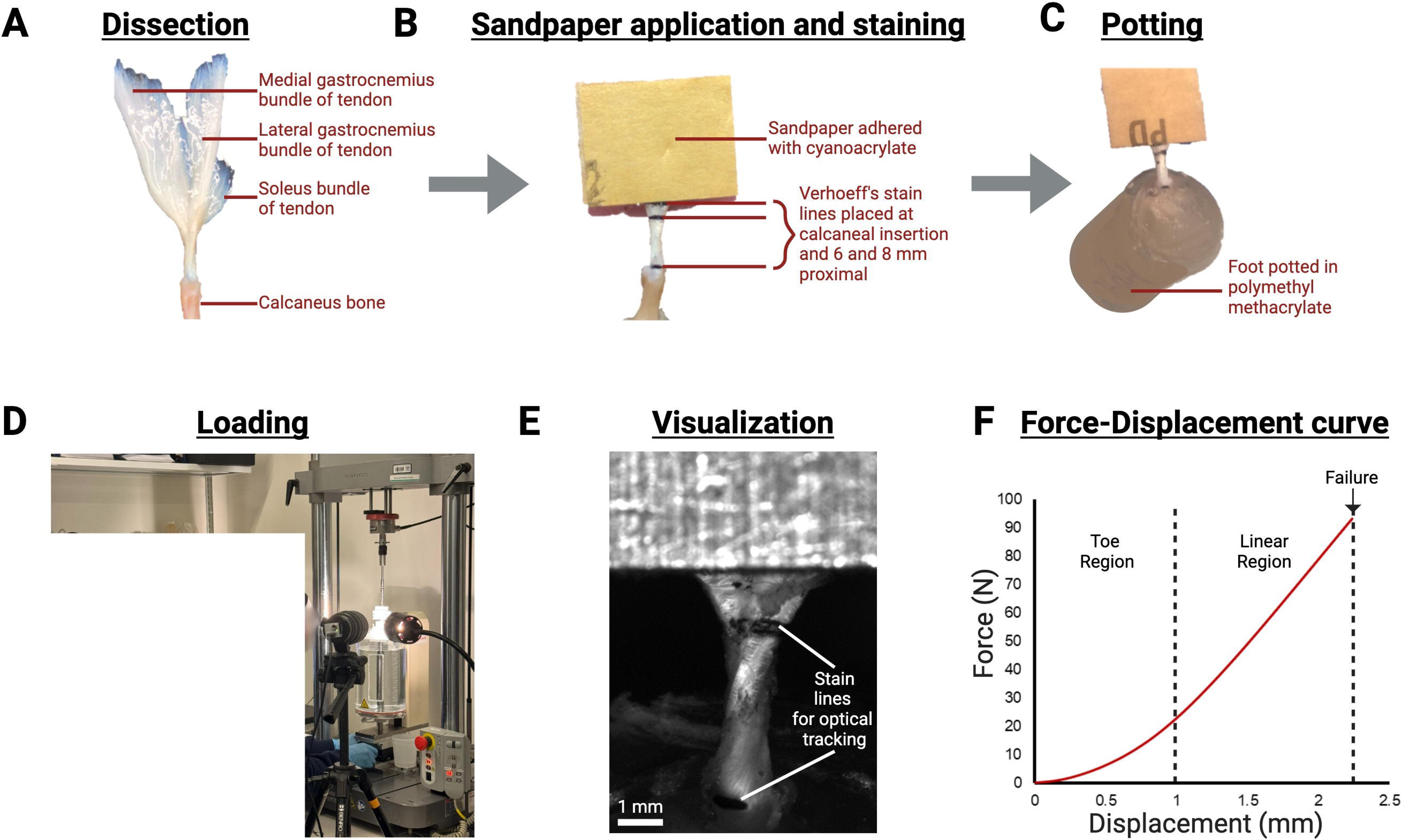

On the day of the module, instructors gave a tour of the PCMD Biomechanics Core, specifically pointing out the uniaxial tensile testing instruments. Next, instructors introduced students to approaches for biomechanical testing of orthopaedic tissues and viscoelastic mechanics. Pairs of students were given a potted foot-tendon complex and told the corresponding rat’s sex and weight. As a group, students hypothesized which tendons would have the highest failure load. Next, with close supervision from instructors, students loaded the potted foot-tendon complex into custom grips, preloaded tendons to 0.1 N, and ran a uniaxial tensile test (ElectroPlus 3000, Natwick, MA) (Figure 2D). The testing protocol included preconditioning (10 cycles from 0.5 to 1.5% strain) followed by quasi-static ramp to failure (0.3% strain per second). Images were recorded (Basler, Exton, PA) and displayed on computer monitors during the tests to ensure all students could see and to demonstrate the concept of optical strain tracking (Figure 2E-F). After tendon testing, instructors led a discussion reviewing the group’s hypotheses and variables that affect tendon mechanical integrity.

### 2.5 Microcomputed tomography (MicroCT) activity

The learning goals of the microcomputed tomography (microCT) activity were to understand how radiography can be used to analyze trabecular bone microstructure and distinguish between healthy and diseased bone.

Prior to the module, instructors scanned tibiae from healthy and ovariectomized rats using the Scanco µCT 45 desktop microCT scanner (Scanco Medical AG, Brüttisellen, Switzerland). Ovariectomy surgery depletes estrogen levels and induces bone loss, serving as a model of osteoporosis.^43^ Scans were captured prior to the first iteration of Learning on a Limb, then re-used for future iterations.

On the day of the module, instructors provided an overview of the ovariectomized rat model, images of healthy versus osteoporotic rat bone, and the technical background of microCT imaging (Figure 3A). The instructors also explained how trabecular bone microstructure can be quantified and described common trabecular microstructure parameters (*e.g.,* bone volume fraction, trabecular thickness).^44^ Next, students were guided through the Scanco microCT image analysis interface and learned to contour regions of interest in the pre-scanned rat tibiae. Each student contoured approximately 10 2D slices in the proximal tibiae of either a healthy or osteoporotic bone scan (Figure 3B-C), then ran a trabecular microstructure analysis. Students were blinded to the type of sample they analyzed; therefore, instructors could challenge them to collaboratively determine which samples were healthy and which were osteoporotic. Lastly, students toured the microCT instruments in the PCMD MicroCT Core and learned how samples are prepared for microCT analysis.

**Figure 3.**
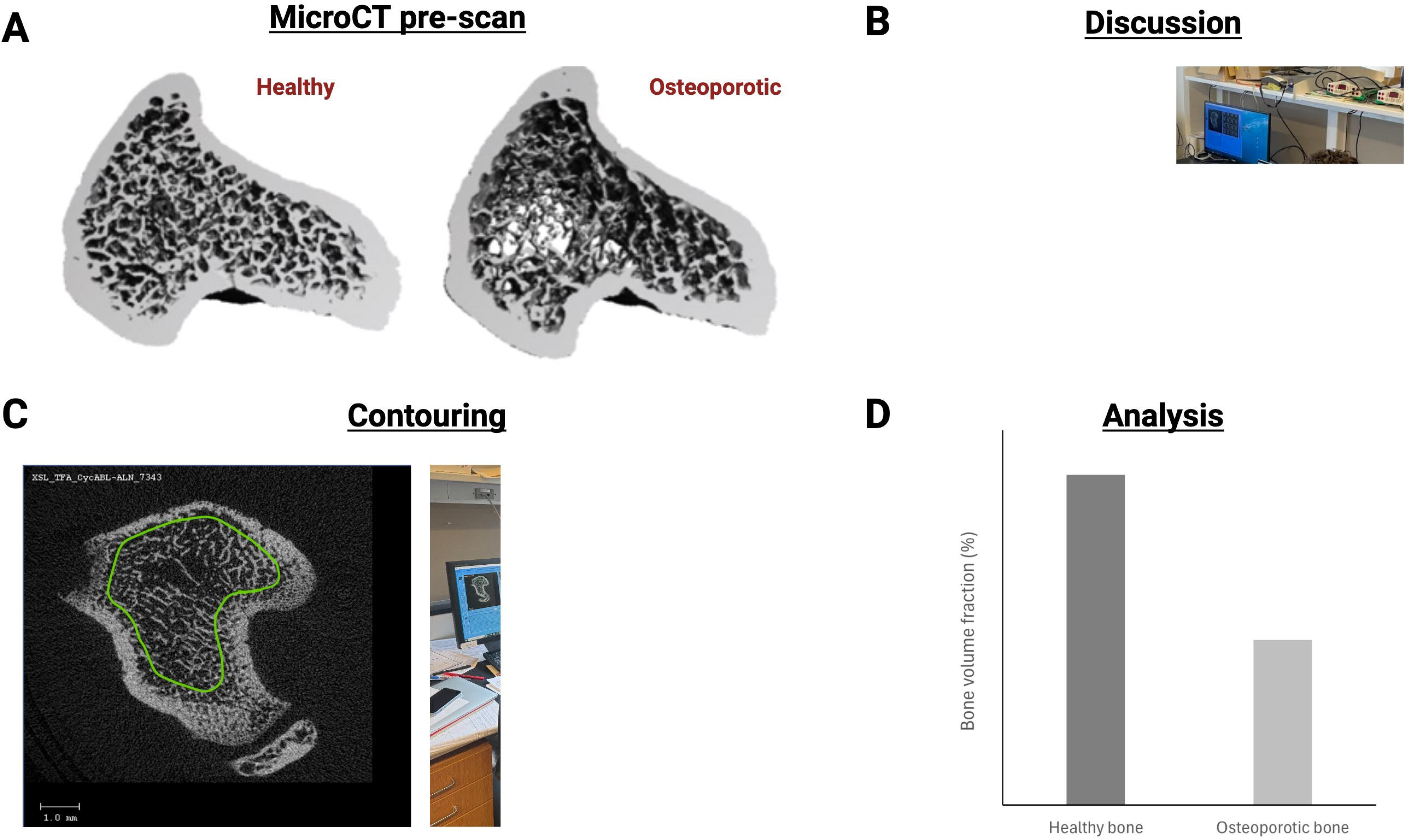
Students contoured healthy and osteoporotic rat tibiae. **(A)** 3D reconstructions of healthy and osteoporotic proximal rat tibiae. **(B)** Students learned about models of osteoporosis and microcomputed tomography (microCT). **(C)** During the activity, students contoured 2D regions of interest in the proximal tibiae, with instructor assistance. **(D)** Pre-compiled data for bone volume fraction, which students used to collaboratively identify differences between healthy and osteoporotic samples.

### 2.6 Cell culture activity

The learning goals for the cell culture activity were to understand techniques for culturing and assaying bone marrow mesenchymal stromal cells (BMSCs).

Prior to the module, BMSCs were isolated from mouse tibiae and cultured for 7-10 days. Tibiae from four mice were dissected and cleaned, then a 23G needle was used to puncture a hole in both ends or the epiphyses of each bone. Next, a hole was punctured into a 0.5 mL centrifuge tube using a 20G needle (Figure 4A). The punctured tibiae were placed in the smaller tube, and the 0.5 mL tube was nested into a larger 1.5 mL centrifuge tube (Figure 4B). This setup was centrifuged at 10,000 RCF for 3 sec to collect BMSCs. Cell pellets were resuspended and cultured in growth medium (α-Minimal Essential Media supplemented with 15% fetal bovine serum, 100 U/mL Penicillin-Streptomycin, and 0.1% 2-mercaptoethanol) for 7-10 days. Cells were seeded at either 0.5 million or 3 million cells per flask to demonstrate the visual appearance of different seeding densities. One flask was prepared for each student; however, students could be placed into small groups of 2-3 to reduce the number of flasks needed.

**Figure 4.**
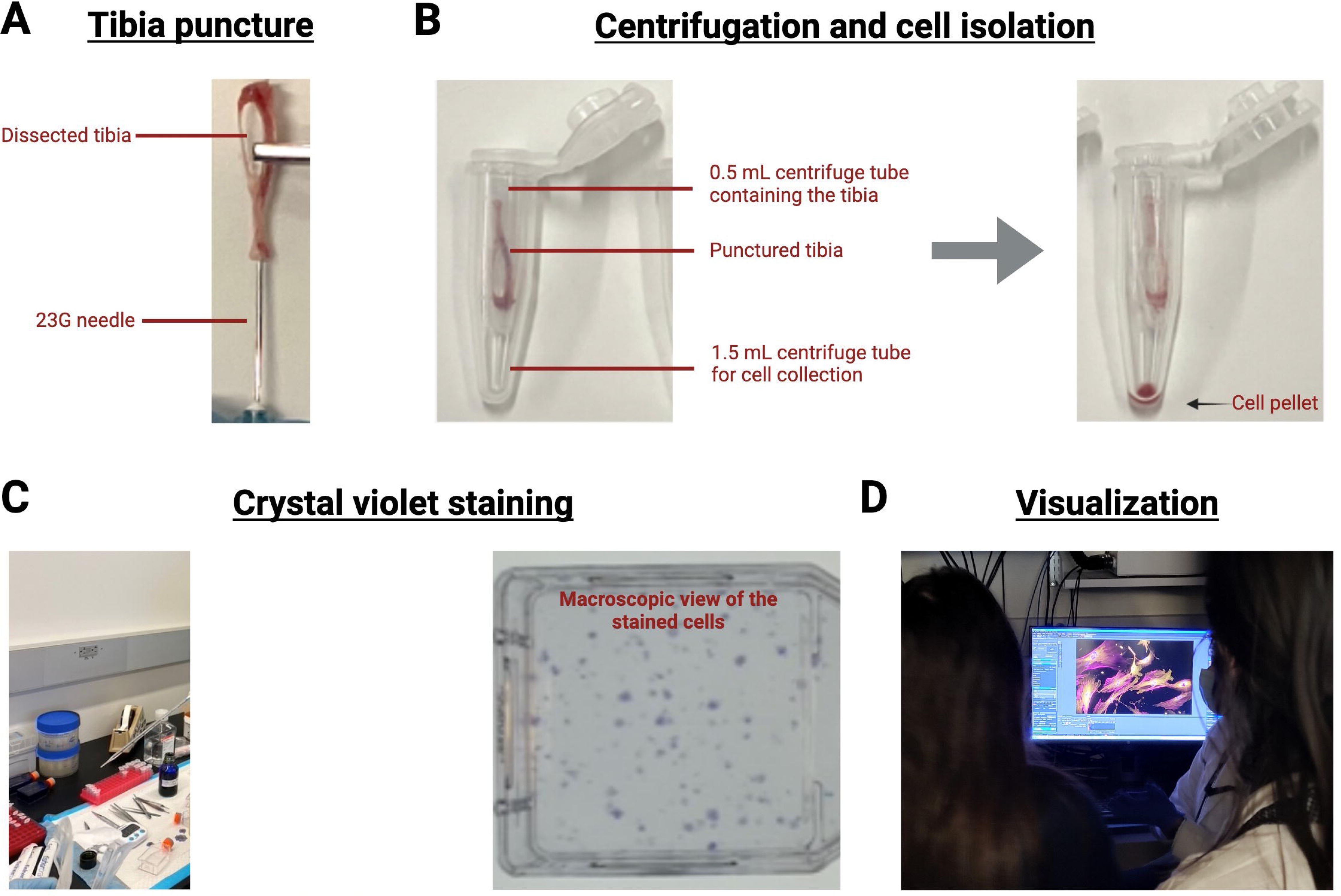
Students cultured mouse bone marrow mesenchymal stromal cells (BMSCs). **(A)** Tibiae were punctured with a 23G needle. **(B)** A 20G needle was used to puncture a hole in a 0.2mL tube. Punctured tibiae were placed in the punctured tube, the punctured 0.2 mL tube was nested within a larger 1.5mL tube, and tubes were centrifuged. The cell pellet was collected at the bottom of the 1.5 mL tube. **(C)** During the activity, students stained BMSCs with crystal violet and visualized cell colonies, with instructor assistance. Each spot stained purple or violet in the 25 cm^2^ cell culture flask is a cell colony. **(D)** Instructors displayed crystal violet-stained BMSCs under a microscope.

On the day of the module, students worked with instructors to isolate BMSCs as described above (Figure 4C). Student-isolated BMSCs were incubated, but students used prepared BMSC flasks to learn sterile tissue culture technique in the biosafety cabinet. They sterilely performed a crystal violet stain to visualize cells cultured in a dish and to count the number of cell colonies.^45^ To conduct the assay, students washed prepared BMSC flasks with 1X PBS twice, then, added 3% crystal violet (w/v) in methanol to the flasks for 15 minutes. Following the 15 minutes incubation, excess crystal violet was recycled and the cells were washed with running tap water (Figure 4D). Instructors guided students in their observations of BMSC colonies by naked eye and under a microscope.

### 2.7 Histology activity

The learning goals of the histology activity were to understand how orthopaedic tissues are prepared for imaging and how to distinguish different orthopaedic tissues from one another.

Prior to the module, rat Achilles tendons and rabbit intervertebral discs (IVDs) were isolated and processed for paraffin histology (Figure 5A). Many paraffin sections were captured from each tissue prior to the first iteration of Learning on a Limb, and stored at room temperature for future iterations of the module.

**Figure 5.**
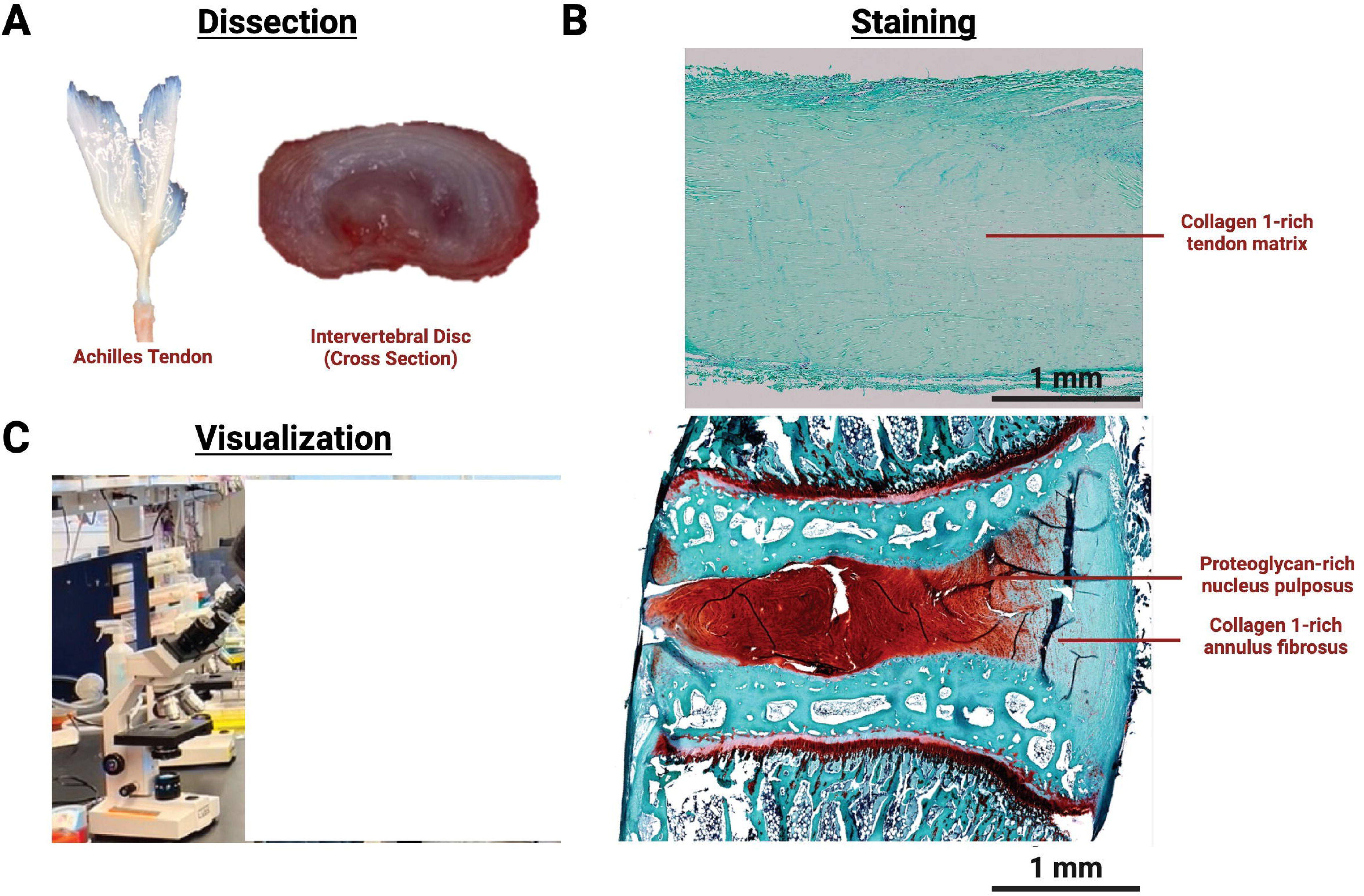
Students histologically analyzed intervertebral disc (IVD) and tendons. **(A)** Prior to the activity, Achilles tendon samples from rats and IVD samples from rabbits and were processed for paraffin sectioning. **(B)** During the activity, students stained sections with safranin O/fast green, with instructor assistance. **(C)** Instructors guided students through visualization of key anatomical features of the tendon and IVD.

On the day of the module students a toured the PCMD Histology Core, with specific attention to the embedding and sectioning equipment. Next, each student was given either a slide section of a rat tendon or a rabbit IVD. Students labeled samples with their names and were guided students through an abridged protocol for safranin O/fast green staining (Table 1). The initial xylene, ethanol (EtOH), and deionized (DI) water incubations were completed prior to students arriving so the stain could be completed in the allotted time. Deparaffinized and rehydrated slides were kept in deionized water until students arrived.

**Table 1.**
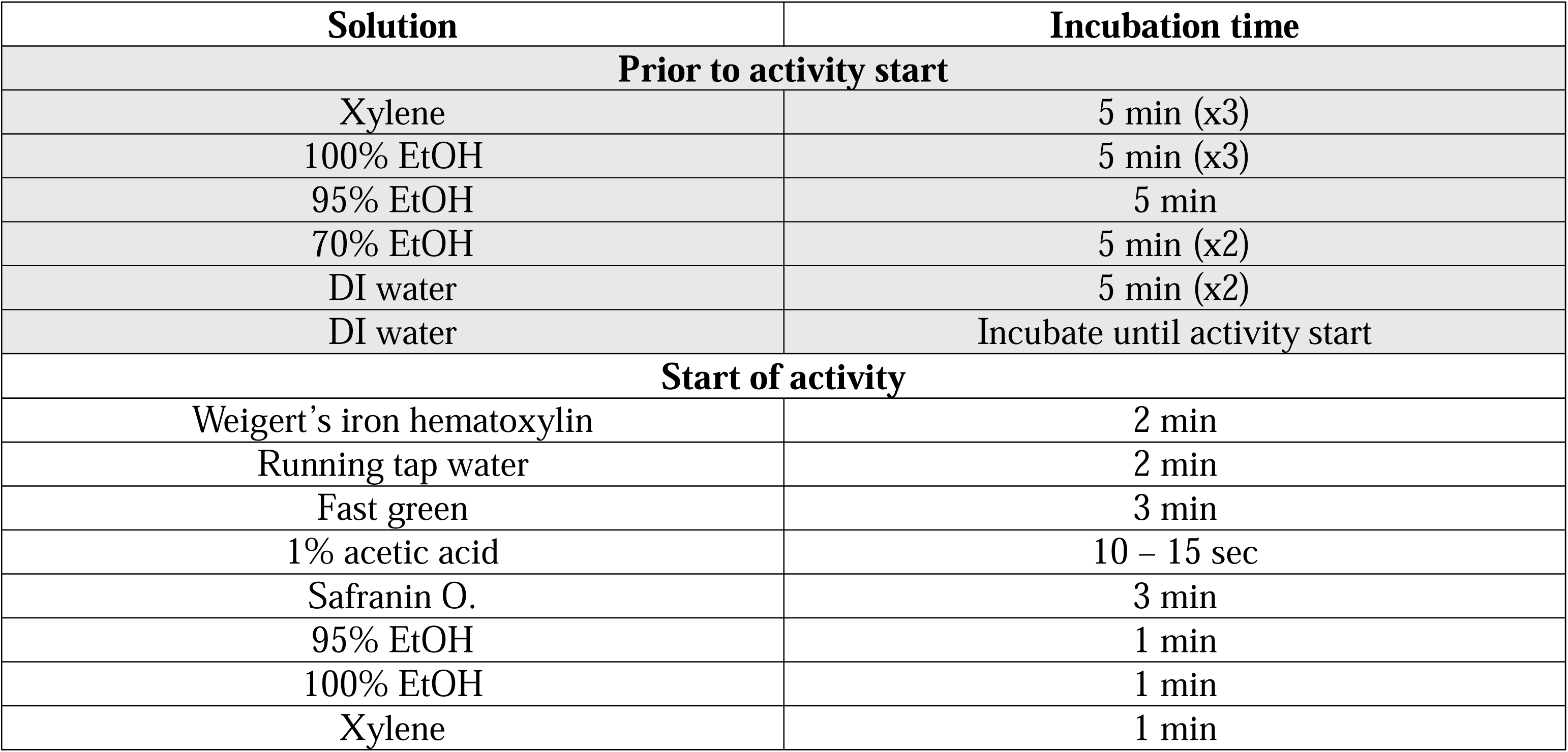
Abridged protocol for safranin O/fast green staining. Gray rows indicate steps completed prior to activity. White rows indicate steps completed with students. EtOH = ethanol.

| Solution | Incubation time |
| --- | --- |
| **Prior to activity start** |  |
| Xylene | 5 min (x3) |
| 100% EtOH | 5 min (x3) |
| 95% EtOH | 5 min |
| 70% EtOH | 5 min (x2) |
| DI water | 5 min (x2) |
| DI water | Incubate until activity start |
| **Start of activity** |  |
| Weigert's iron hematoxylin | 2 min |
| Running tap water | 2 min |
| Fast green | 3 min |
| 1% acetic acid | 10 – 15 sec |
| Safranin O. | 3 min |
| 95% EtOH | 1 min |
| 100% EtOH | 1 min |
| Xylene | 1 min |

After staining, students coverslipped their samples and viewed them under a Celestron^TM^ CB2000CF Compound Microscope (Celestron, Torrance, CA, USA) (Figure 5B). Instructors asked students guiding questions to highlight the differences between tendons and IVDs and the differences between the nucleus pulposus and annulus fibrosus IVD regions (Figure 5C).

### 2.8 Post-activity exercises

During the post-activity exercises, students discussed what they learned and participated in a speed-networking session. Similar to the pre-activity exercise, we employed active learning, which encouraged students to think critically about the techniques they learned could be used for additional research. During the speed networking session, student activity groups were matched with 2-3 Learning on a Limb instructors to learn how and why they pursued careers in orthopaedic research. Student activity groups met with two sets of Learning on a Limb instructors for approximately 15 minutes each.

### 2.9 Learning module evaluation

To evaluate student learning, we developed pre/post-tests consisting of eight multiple choice test questions (TQs) that assessed student comprehension of how biomechanical testing, microCT, cell culture, and histology are applied to orthopaedic research (Appendix A). Pre/post-tests were administered to the 2023 and 2024 cohorts. This test length was similar to published biomedical engineering education modules^23,24,26,39,40^ and allowed us to test two concepts for each technique. Pre-tests were administered at the beginning of the pre-activity exercises and post-tests at the end of the post-activity exercises. Students did not have the opportunity to review their pre-test results to avoid the possibility of students simply recalling correct test answers.

To evaluate student attitudes towards orthopaedic research and biomedical engineering, we developed a post-survey (Appendix B). The survey questions (SQs) were designed to evaluate how the outreach module impacted student attitudes towards orthopaedics and biomedical engineering. For each SQ, students were asked to rate the influence Learning on a Limb had: “No Influence”, “Slight Influence”, “Definite Influence”, or “Major Influence”. Post-surveys were administered at the end of the post-activity exercises (short-term post-survey) and at 1 year after participating in Learning on a Limb (long-term post-survey). Short-term post-surveys were administered to the 2023 and 2024 cohorts, while long-term post-surveys were administered to the 2022 and 2023 cohorts.

Instructors anonymized pre/post-tests and post-surveys prior to grading and data analysis. Statistical analysis was conducted using GraphPad Prism® software version 10 (GraphPad Software, San Diego, CA, USA). To determine significant learning gains, pre/post-test scores were compared using a paired Student’s t-test (α = 0.05).

## 3.0 Results

### 3.1 Student demographic information

During three years of implementing Learning on a Limb, we recruited 32 students from several high schools in the Greater Philadelphia Area (N = 32). An approximately even number of 10^th^, 11^th^, and 12^th^ grade students were recruited to participate in Learning on a Limb (Figure 6A). Students identified as Asian (53.13%), Black (25.00%), LatinX (9.38%), or Other (3.13%); thus, we successfully recruited diverse students to complete Learning on a Limb (Figure 6B). Additionally, 71.88% of students identified as Female, indicating that most students who completed Learning on a Limb were gender minorities in STEM (Figure 6C).

**Figure 6.**
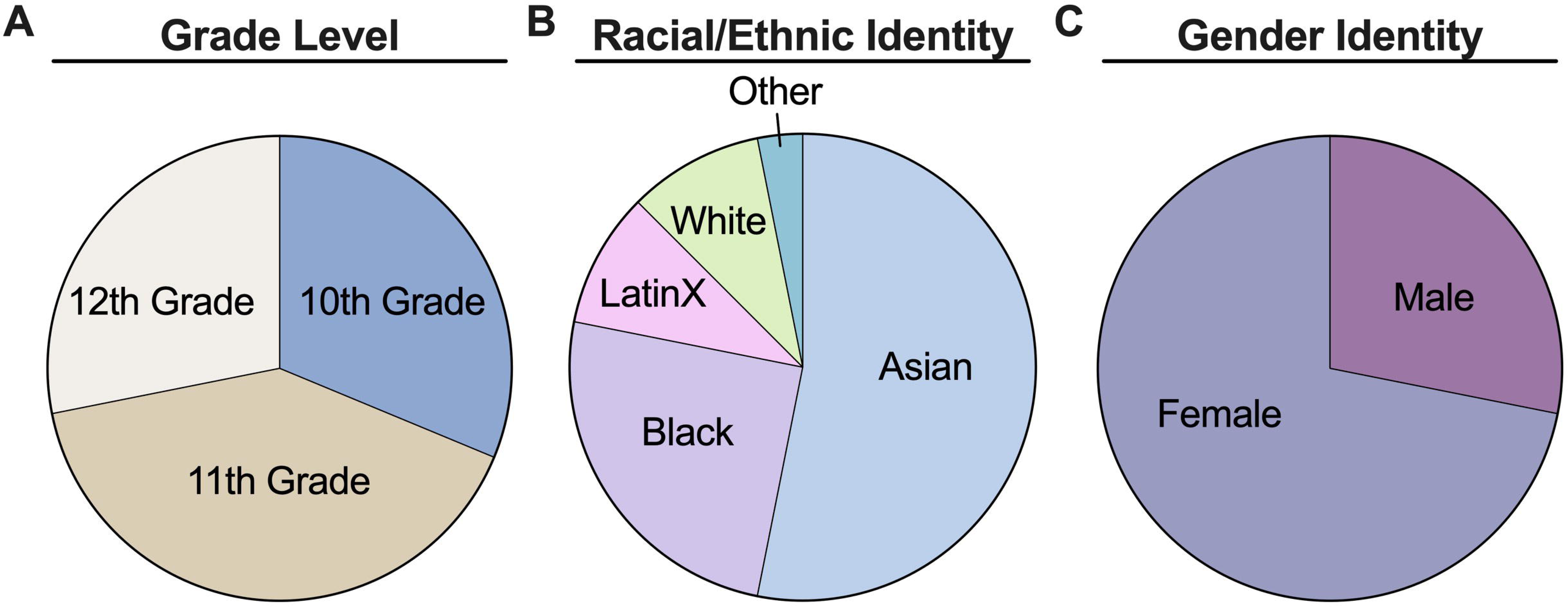
Students belonged to various grade levels and most identified as racial/ethnic or gender minorities in orthopaedics. **(A)** Breakdown of high school grade. **(B)** Breakdown of racial/ethnic identity data. **(C)** Breakdown of gender identity data. N = 32 students.

### 3.2 Pre/post-test knowledge assessments

Pooled pre/post-test assessments were scored for two cohorts of high school students to evaluate knowledge gains from Learning on a Limb (N = 17). Instructors designed the pre/post-tests to test concepts from each hands-on activity (Appendix A). Results showed that students experienced significant learning gains with average test scores increasing from 27.2% on the pre-test to 78.7% on the post-test (Figure 7A). In addition to significant increases in overall score, we found that the percentage of students who answered each test question (TQ) correctly was greater in the post-test than the pre-test (Figure 7B). A very low percentage of students answered most TQs correctly in the pre-test, but a majority answered all TQs correctly in the post-test. Overall, our results indicate that students had very little prior exposure to the content covered in Learning on a Limb and that completing this module significantly increased their knowledge of orthopaedics.

**Figure 7.**
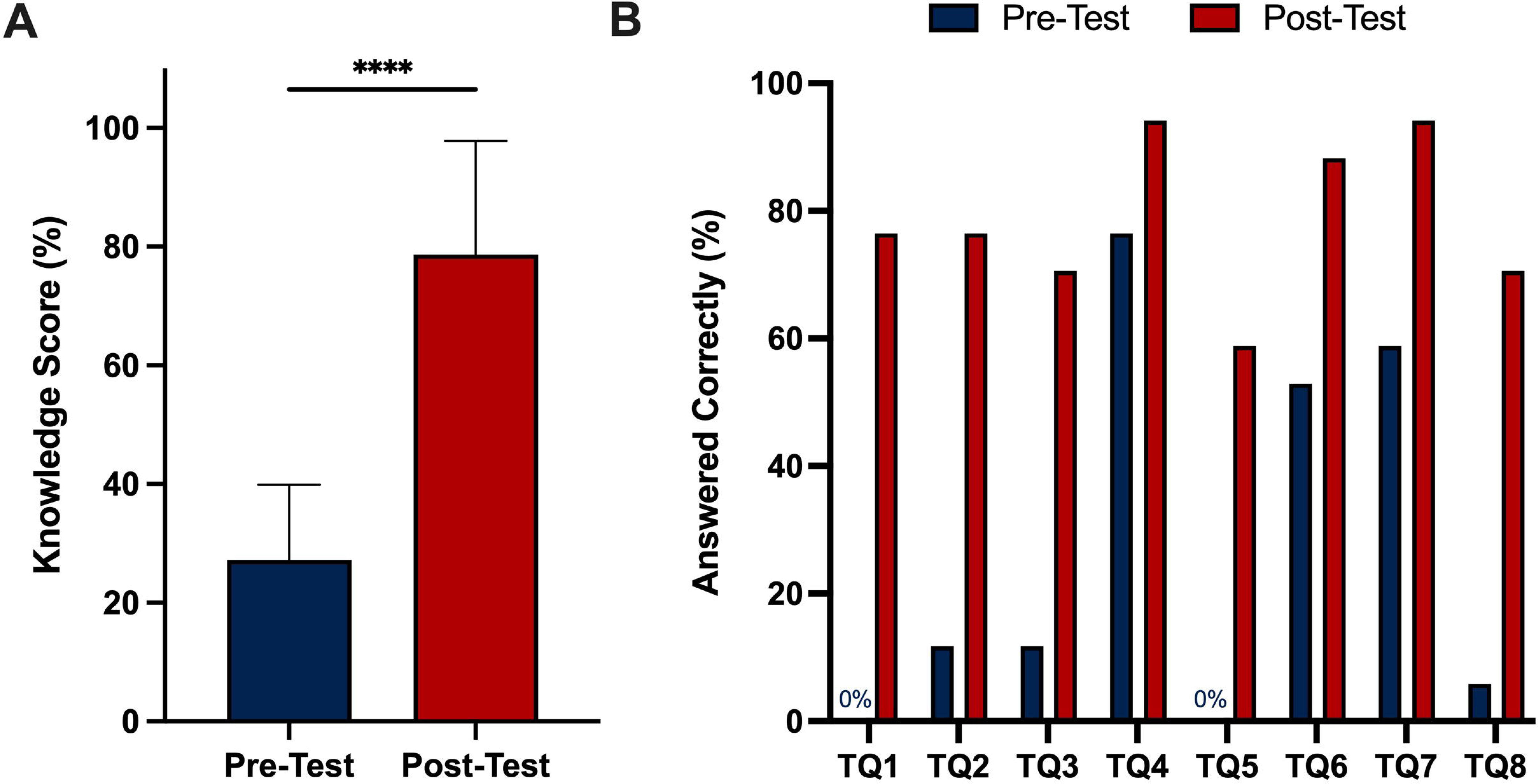
Students showed significant learning gains by participating in Learning on a Limb. **(A)** Average student pre/post-test scores presented as mean ± standard deviation. **(B)** Percentage of students who answered individual questions correctly on pre/post-tests. **** = p<0.0001 using paired. Student’s t-test. N = 17 students.

### 3.3 Short-term post-survey attitudes assessment

Pooled short-term post-survey assessments were collected for two cohorts to evaluate attitude changes from Learning on a Limb (N = 17). Instructors designed the post-survey to evaluate student attitudes towards orthopaedics and biomedical engineering (Appendix B). After completing the module, at least 50% of the students indicated that Learning on a Limb had a “Major” or “Definite” influence on their interest in learning about orthopaedics and biomedical engineering (SQ1, SQ2, SQ4, & SQ5) and pursuing careers in orthopaedics and biomedical engineering (SQ3 & SQ6) (Figure 8). Thus, our results demonstrated that Learning on a Limb is an effective module for sparking student interest in orthopaedic-related research and career opportunities.

**Figure 8.**
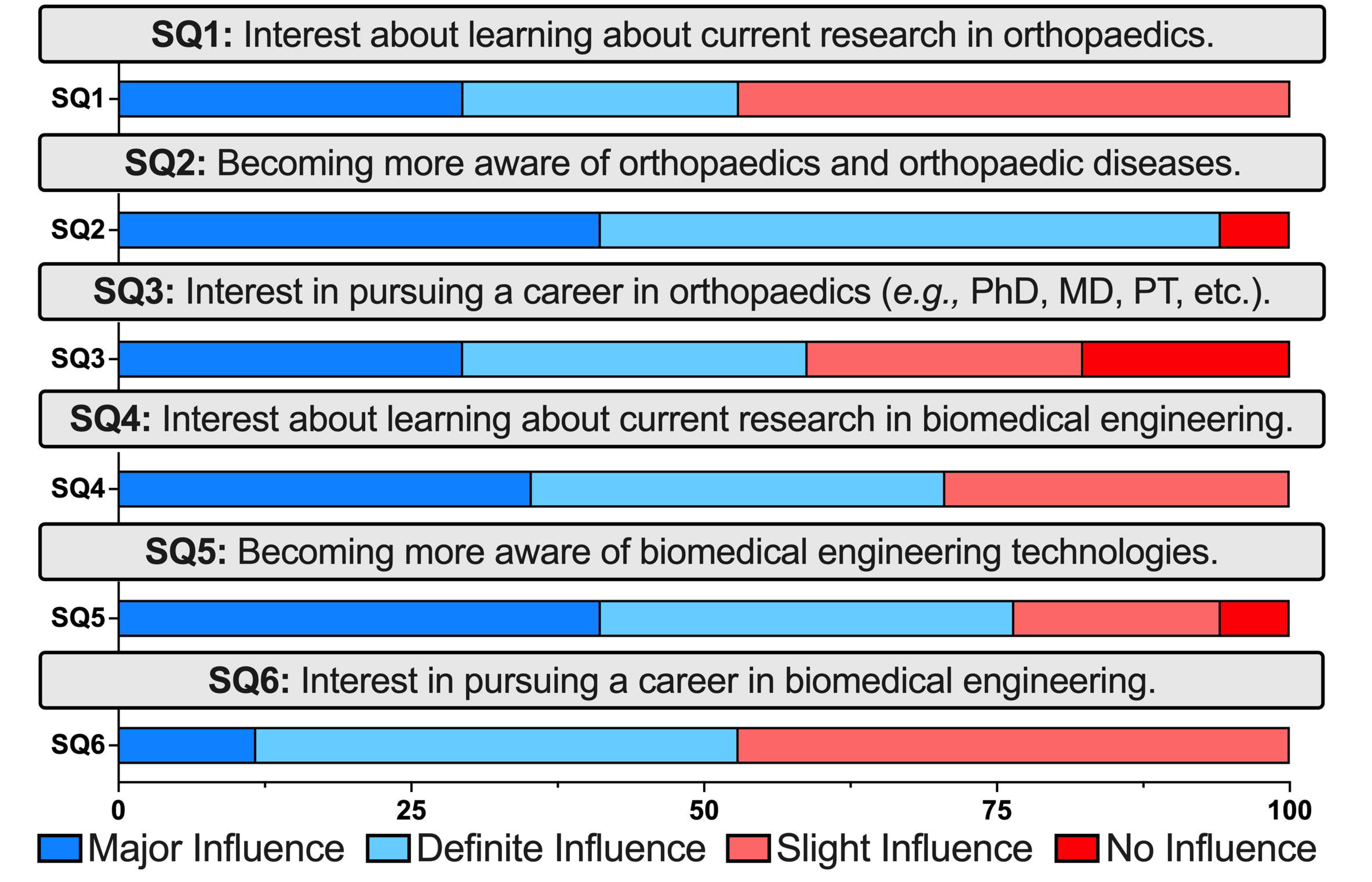
Short-term post-survey showing Learning on a Limb influenced student interest in orthopaedics and biomedical engineering. N = 17 students.

### 3.4 Long-term post-survey attitudes assessment

Learning on a Limb graduates were longitudinally tracked by the Perelman School of Medicine Office of Outreach, Education, and Research (OER) to determine longer-term participation benefits. Among the 32 Learning on a Limb graduates, 71.88% (23/32 = 71.88%) participated in summer research experiences at the University of Pennsylvania and 25.00% (8/32 = 25.0%) completed their research in the McKay Orthopaedic Research Laboratory (Figure 9A-B). These findings demonstrate how orthopaedic outreach modules, like Learning on a Limb, can be an effective pipeline tool for recruiting talented high school students into orthopaedic research.

**Figure 9.**
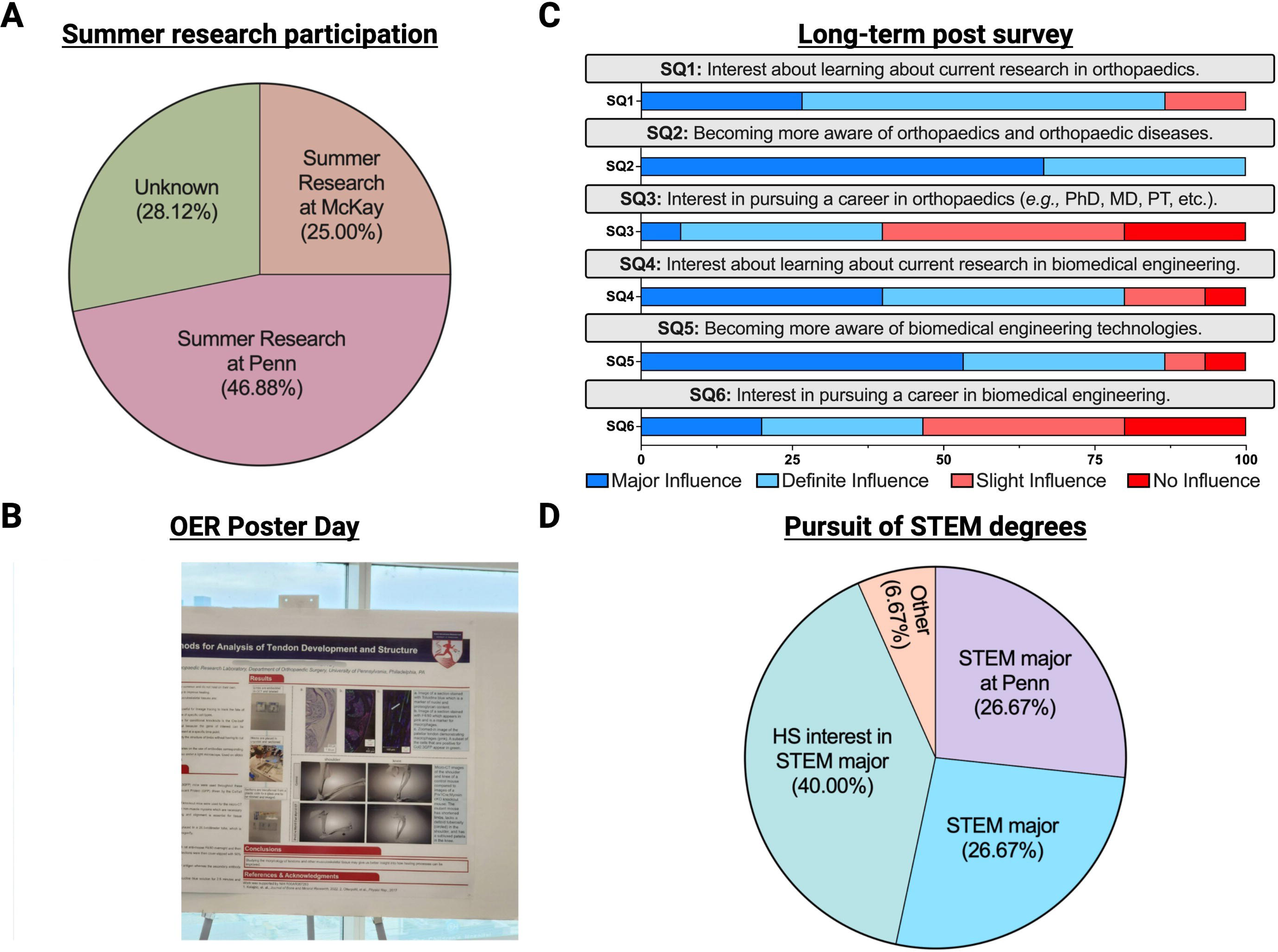
Long-term post-survey showing Learning on a Limb had sustained influence on student interest in orthopaedics and biomedical engineering. **(A)** Breakdown of Learning on a Limb graduates participating in summer research experiences. N = 32 students. **(B)** Learning on a Limb graduate presenting at the University of Pennsylvania Office of Outreach, Education, and Research (OER) Poster Day. **(C)** Long-term post-survey data demonstrating sustained interest in orthopaedics and biomedical engineering after 1 year. N = 15 students. **(D)** Breakdown of Learning on a Limb graduates pursuing STEM majors in college.

We also administered a long-term post-survey to the 2022 and 2023 cohorts at 1 year after participating in Learning on a Limb. Approximately 67% (15/27 = 66.67%) of students completed this long-term post-survey. Greater than 75% of students indicated that Learning on a Limb had a “Major” or “Definite” influence on their interest in learning about orthopaedics research (SQ1 & SQ2) and biomedical engineering (SQ4 and SQ5) (Figure 9C). In addition to retained enthusiasm for orthopaedic and biomedical engineering research, approximately 40% of students indicated that Learning on a Limb had a “Major or “Definite” influence on their interest in pursuing careers in orthopaedics (SQ3) or biomedical engineering (SQ6). These long-term evaluations of our cohorts indicate that Learning on a Limb had a sustained influence on participants and suggest that similar programs may enhance participation in orthopaedics.

Qualitative data from the long-term post-survey showed Learning on a Limb had unexpected positive influences on student confidence, learning, and peer engagement. Almost all participants reported that they shared what they learned during Learning on a Limb with teachers and classmates (13/15 = 86.67%), which shows their confidence with this information. One participant stated, “Personally, I enjoyed being able to have a higher-level discussion about scientific research with my teacher…it was a really interesting conversation where we drew parallels between the research being done in orthopaedics labs to other types of research in the field of biology.” Such testimonials show how students were able to apply the information they learned and analyze connections between orthopaedic research and other biology topics. Another participant shared, “One of my friends is really interested in bioengineering and orthopedic engineering. I was able to teach her about everything I learned during Learning on a Limb. She also attends Penn and is now looking for a lab to work in.” This peer sharing shows how participants can become ambassadors of orthopaedic research, and how Learning on a Limb can have multiplicative effects on promoting involvement in musculoskeletal research.

The long-term post-survey also longitudinally tracked student college trajectories. Among the participants who completed the 1-year follow up, 26.67% (4/15 = 26.67%) were pursuing STEM majors at the University of Pennsylvania and 26.67% (4/15 = 26.67%) were pursuing STEM majors at other institutions (Figure 9D). Excitingly, all participants who were still in high school students (6/15 = 40%) indicated a strong interest in pursuing a STEM major in college. It is unlikely that participation in Learning on a Limb was solely responsible for these college trajectories, since students reported participating in other high school STEM programs. Rather, these results exemplify how hosting engaging K-12 outreach programs can be part of a progressive pathway for URM students to access opportunities in STEM.

## 4.0 Discussion

We must inspire a diverse next generation of orthopaedic researchers to combat the high socioeconomic burden of musculoskeletal disorders. There are several educational modules that implicitly teach college-level students about orthopaedic research; however, there are no published modules to introduce K-12 students to orthopaedics. Here, we designed, implemented, and evaluated Learning on a Limb, an orthopaedic outreach module that teaches diverse high school students about orthopaedic research.

In this program, we aimed to recruit diverse students, who are considered URM students in STEM,^46^ to participate in Learning on a Limb. We made this choice because orthopaedic research and orthopaedic surgery have demonstrated noticeable gaps in racial, ethnic, and gender diversity.^47–51^ Towards this goal, the McKay Orthopaedic Research Laboratory worked with the Perelman School of Medicine Office of OER to recruit diverse students from local high schools in the Greater Philadelphia Area. Widespread implementation of engaging modules, like Learning on a Limb, by orthopaedic departments has the capacity to increase diversity in the future orthopaedic research workforce. These efforts can complement other programs like the Perry Initiative, which empowers women to pursue careers in orthopaedics.^52^

Learning on a Limb participants experienced significant learning gains by completing the module. Almost all concepts were new to students, as indicated by their low pre-test scores. TQ4, which tested how osteoporosis impacts bone mineral density, was the exception and implies that students understood this concept prior to the module. Regardless, students experienced learning gains of 51 points, which was similar to freshman engineering students^53,54^ and middle school students ^39,40^ who completed analagous hands-on engineering activities. Qualitative results from the long-term post-survey suggest that students learned at higher levels than tested. Pre/post-tests asked students to remember facts and understand basic concepts, which are lower-level objectives by Bloom’s Taxonomy.^55^ Unexpectedly, students described their ability to apply this information to new situations and analyze connections between orthopaedics and other research fields, which displays higher-level learning according to Bloom’s Taxonomy. Overall, these significant learning gains demonstrated that Learning on a Limb is an effective module to teach students about orthopaedic research.

Post-survey data showed that Learning on a Limb sparked sustained student interest in orthopaedic research and enhanced student confidence. Most students indicated that participating in Learning on a Limb had a “Major” or “Definite” influence on their interest in orthopaedics, biomedical engineering, and related careers. After completing Learning on a Limb, most participants also went on to participate in summer research experiences at the University of Pennsylvania and pursue STEM majors in college. These results demonstrate how educational modules, like Learning on a Limb, can provide high school students with early exposure to ongoing research, which is a key step to becoming a successful researcher.^56^ Moreover, they can serve as an effective pipeline for connecting URM students to orthopaedics departments and universities, where they can continue finding opportunities to advance their STEM careers. Sharing their experiences with their teachers and peers also indicates that Learning on a Limb helped enhance scientific confidence, which is extremely important for young URM students who report having lower confidence in STEM.^57–59^

Hosting Learning on a Limb synergistically benefitted the participants and the instructors. The Learning on a Limb instructors were a diverse team of principal investigators, postdoctoral research fellows, graduate students, and research scientists at the McKay Orthopaedic Research Laboratory. We aimed to recruit diverse instructors from the department because students are more likely to persist in STEM when they are taught by instructors of the same race and gender.^60^ Instructors expressed a strong sense of satisfaction with leading Learning on a Limb because it provided them with an opportunity to “pay it forward” to the next generation of students. Instructors also reported that participating in Learning on a Limb helped create a sense of community and comradery within the department. This enhanced sense of community is exemplified by this manuscript, which has authors from seven different groups in the McKay Orthopaedic Research Laboratory.

Orthopaedics departments around the world can host Learning on a Limb, as described, or with modifications. For recruitment, we recommend working with on-campus offices and centers that have established connections with local high schools (*e.g.,* Perelman School of Medicine Office of OER). If establishing new connections, it is helpful to connect these contacts with established on-campus offices and centers to facilitate longitudinal tracking. Orthopaedic departments should also make efforts to reduce barriers to participating in the program (*e.g.,* providing lunch and transportation to participating students). Specific activities can be modified based on institutional facilities, research interest areas, and allotted timing for each activity. If reducing the number activities, instructors could include more conditions into the remaining activities (*e.g.,* young vs. old, male vs. female, diseased vs. healthy), then guide students through quantitative analyses. For example, students could quantify how different factors affect (1) tendon tensile properties, (2) trabecular microstructure, (3) colony forming units, or (4) histological grading. Orthopaedics departments can take advantage of the low-cost and adaptable nature of this module to provide URM students with tangible benefits.

## 5.0 Conclusion

This study designed, implemented, and evaluated Learning on a Limb, an orthopaedic outreach module to teach diverse high school students about orthopaedic research. Over the course of this 4-hr module, students completed hands-on activities to learn how biomechanical testing, microCT, cell culture, and histology are used in orthopaedic research. We recruited 32 high school students from the Greater Philadelphia Area to participate in Learning on a Limb over the past three years, most of which identified as racial/ethnic or gender minorities in orthopaedic research. Overall, we found that completing Learning on a Limb resulted in significant learning gains and increased interest in orthopaedic research. Learning on a Limb also served as an effective pipeline for recruiting diverse students to research at the University of Pennsylvania. In addition to student benefits, the instructors benefited by having the opportunity to “pay it forward” to the next generation of students and build community within their department. Broad deployment of this module by orthopaedics departments would synergistically inspire diverse students to study orthopaedics, strengthen community within orthopaedics departments, and promote innovations in orthopaedic research.

## 6.0 Acknowledgements

The authors thank Dr. Daphney R. Chery for the original conception of this idea and Tarence Smith for help recruiting participants for Learning on a Limb. The authors also thank Dr. Sereen S.F. Assi, Rebecca Betts, Dr. Jonathan Blank, Dr. Kevin G. Burt, Mary Kate Evans, Sanjana Hemdev, Dr. Yongqiang Vincent Jin, Talayah Johnson, Dr. Annemarie Lang, Dr. Liane Miller, Felicia Pinto, Dr. Karthik Rajagopal, Dr. Jaime Santillan, Elizabeth Seidl, Dr. Ke Song, Nat Thurlow, Stephanie Weiss, and Dr. Xiaoyu Xu for assisting with Learning on a Limb activities. Figures 1 – 5 and 9 were created using Biorender.com.

This work was supported by the Penn Center for Musculoskeletal Disorders [NIH/NIAMS P30-AR069691], the Penn Achilles Tendinopathy Center for Research Translation [NIH/NIAMS P50-AR080581], the University of Pennsylvania Institutional Research and Academic Career Development Award (IRACDA) [NIH/NIGMS K12GM081259], the Department of Veterans Affairs Rehabilitation Research and Development Service [IK2 RX003118], the Cali/Weldon Professorship, and the Training in Musculoskeletal Research Grant [NIH/NIAMS 5-T32-AR007132].

## Appendix A – Pre/post-test questions assessing knowledge of orthopaedics

1. Which of the following mechanical tests can determine the tensile stiffness of a tendon?

a. Repeatedly twisting the tendon
b. **Pulling the tendon apart so it is elongated**
c. Pushing the tendon together so it is shortened
d. Repeatedly pulling and pushing the tendon, so it is elongated and shortened
e. I don’t know
2. How would the mechanical properties of a healthy tendon be different from a diseased tendon?

a. **Healthy tendons would be stiffer**
b. Diseased tendons would be stiffer
c. Healthy and diseased tendons would have the same stiffness
d. I don’t know
3. Micro-computed tomography (Micro-CT) can be used to analyze which tissue type?

a. Tendons
b. Ligaments
c. Muscle
d. **Bones**
e. I don’t know
4. How would the bone density of a healthy skeleton be different from a skeleton with osteoporosis?

a. **Healthy skeletons would have higher mineral density**
b. Skeletons with osteoporosis would have higher mineral density
c. Healthy skeletons and skeletons with osteoporosis would have the same mineral density
d. I don’t know
5. Which assay can be used to identify mesenchymal progenitor cells?

a. Polymerase Chain Reaction Assay
b. **Colony Forming Unit Assay**
c. Western Blot Assay
d. Bicinchoninic Acid (BCA) Assay
e. I don’t know
6. Where can you find mesenchymal progenitor cells in the body?

a. Skin
b. Eyes
c. **Bone**
d. Muscle
e. I don’t know
7. What are the steps a researcher would use to prepare a tissue for histology?

a. **Section, stain, image with a microscope**
b. Stain, section, image with a microscope
c. Section, image with a microscope, stain
d. Stain, image with a microscope, section
e. I don’t know
8. Which tissue would stain positively for Safranin O?

a. Tendon
b. Muscle
c. Annulus fibrosus
d. **Nucleus Pulposus**
e. I don’t know

## Appendix B - Post-survey questions assessing attitudes towards orthopaedic research

On a scale from 1-4, how much of an influence has Learning on a Limb had on you in the following areas? (Where 1 means “No Influence”, 2 means “Slight Influence”, 3 means “Definite Influence”, and 4 means “Major Influence”.)

1. Interest about learning about current research in orthopaedics
2. Becoming more aware of orthopaedics and orthopaedic diseases
3. Interest in pursuing a career in orthopaedics (e.g., PhD, MD, Physical Therapy, etc.):
4. Interest about learning about current research in biomedical engineering.
5. Becoming more aware of biomedical engineering technologies.
6. Interest in pursuing a career in biomedical engineering.

Please describe the most memorable part of your Learning on a Limb experience.

---

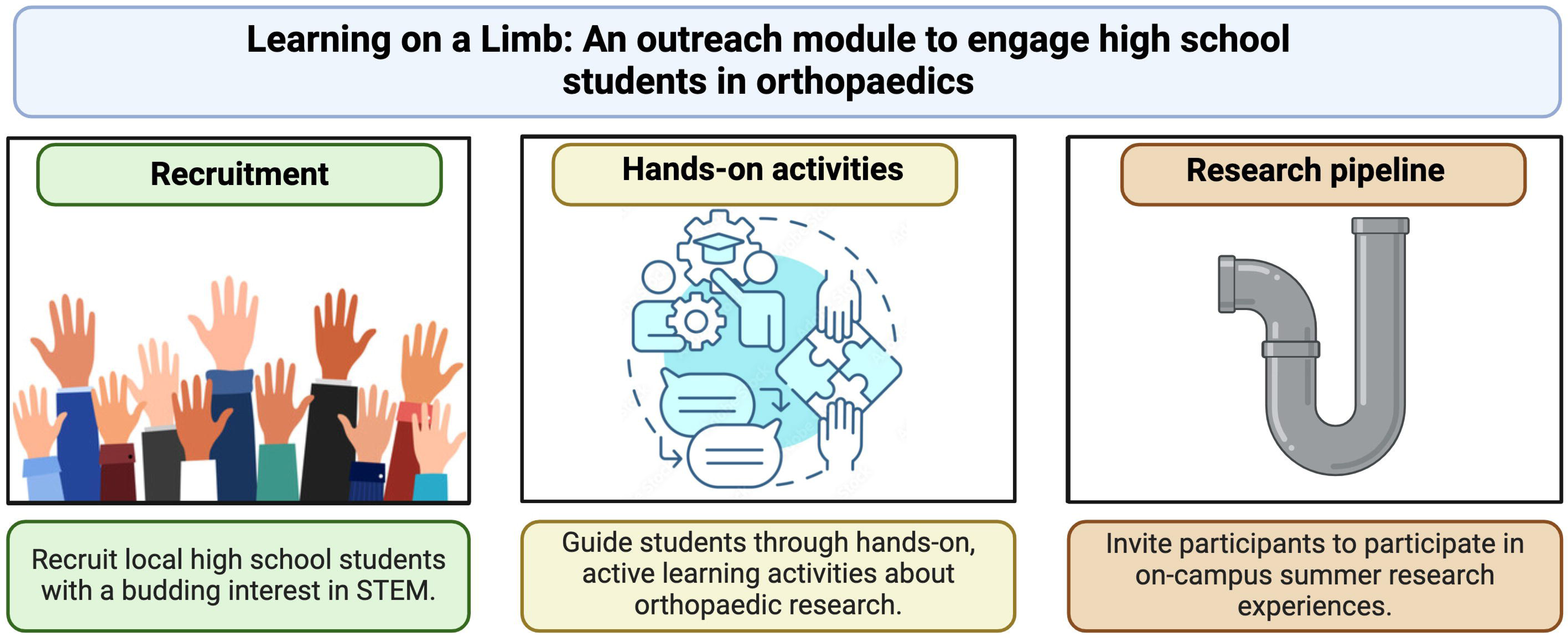

